# Dynamics of repair and regeneration of adult zebrafish respiratory gill tissue after cryoinjury

**DOI:** 10.1101/2021.05.27.445469

**Authors:** Marie-Christine Ramel, Fränze Progatzky, Anna Rydlova, Madina Wane, Jürgen Schymeinsky, Cara Williams, Birgit Jung, Jonathan Lamb, Matthew J Thomas, Laurence Bugeon, Margaret J. Dallman

## Abstract

The study of respiratory tissue damage and repair is critical to understand not only the consequences of respiratory tissue exposure to infectious agents, irritants and toxic chemicals, but also to comprehend the pathogenesis of chronic inflammatory lung diseases. To gain further insights into these processes, we developed a gill cryoinjury model in the adult zebrafish. Time course analysis showed that cryoinjury of the gills triggered an inflammatory response, extensive cell death and collagen deposition at the site of injury. However, the inflammation was rapidly resolved, collagen accumulation dissipated and by 3 weeks after injury the affected gill tissue had begun to regenerate. RNA seq analysis of cryoinjured gills, combined with a comparison of zebrafish heart cryoinjury and caudal fin resection datasets, highlighted the differences and similarities of the transcriptional programmes deployed in response to injury in these three zebrafish models. Comparative RNA seq analysis of cryoinjured zebrafish gills with mouse pulmonary fibrosis datasets also identified target genes, including the understudied FIBIN, as differentially expressed in the two species. Further mining, including of human datasets, suggests that FIBIN may contribute to the successful resolution of tissue damage without fibrosis.

## Introduction

There are limited treatment options for people suffering from substantive lung damage. As a consequence, chronic respiratory diseases are a leading cause of mortality and morbidity; it has been estimated that in 2017 half a billion people worldwide were affected by a chronic respiratory disease (GBD Chronic Respiratory Disease report, 2020). Bacterial/viral infections, inhalation of toxins such as cigarette smoke, or exposure to environmental particulates and pollutants such as diesel fumes may all cause extensive damage to respiratory tissue and this is accompanied by unresolved chronic inflammation and consequent loss of lung function [1-3]. A recent striking example is the current Covid-19 pandemic in which infection with the SARS-CoV-2 virus can, in severe cases, lead to an overactive pulmonary immune response and lung alveolar damage [4]. Emerging evidence also indicates that a subset of recovering patients also suffer from pulmonary fibrosis after infection [5]. In the absence of suitable cures, there is a crucial need to develop experimental research models that contribute to our understanding of lung injury, inflammation, repair and regeneration.

Studying the biology of diseased lung tissue using human clinical specimens is challenging, with samples being difficult to obtain in the early stages of disease. In addition, control ‘healthy’ samples are highly variable in terms of patients’ age and underlying health conditions. Consequently, not only is a basic comparison of healthy and diseased lung tissue difficult, but intervention studies are also complex. Researchers have therefore turned to animal models to develop an understanding of not only the pathogenesis and progression of human lung disease but also the general response of the respiratory tissue after exposure to infectious agents or chemicals/irritants. Mouse studies have been critical in understanding many aspects of human lung tissue responses and pathologies with other mammalian models such as the guinea pig, hamster, rabbit and ferret increasingly used because of more comparable lung anatomy, immune responses and/or reaction to stimuli, irritants and pathogens [6-8].

Although they do not possess lungs, several studies including our own, have recently highlighted the potential of the adult zebrafish (*Danio rerio*) as an alternative model system for respiratory tissue investigations of inflammation and repair [9-12]. In the adult zebrafish, the respiratory organs are the gills which are located in branchial chambers on each side of the head under the protection of a bony flap, the operculum. Zebrafish possess 4 pairs of gill arches which form the base for gill filaments (see Fig. 1K and Fig. S1 for anatomy). Gill filaments are composed of primary lamellae and secondary lamellae where gas exchange occurs. Gill tissue is easily accessible for manipulation; the operculum can be lifted for the application of inflammatory reagents directly to the gill [10] or can be removed for live imaging or physical manipulation such as resection [11]. As with the alveolar surface of the mammalian lung, zebrafish/teleost gills are covered by a mucosal layer which acts as a first barrier of defence against waterborne pathogens and irritants [13]. The gill tissue is populated by a variety of immune cells of both innate and adaptive immunity [9, 14](Wane, 2021 PhD thesis). Lymphocytes in particular accumulate at the base of the filaments and along the filaments in a structure termed interbranchial lymphoid tissue [15, 16]. We have recently shown that the zebrafish gill immune system can mount an inflammatory response when gill tissue is exposed to cigarette smoke extracts [9] or a viral adjuvant [10] and this response is altered in *il4*/*13* and *il10* zebrafish mutants [17]. How the immune reaction contributes to specific responses of the respiratory tissue to a variety of stimuli, including physical damage and exposure to toxicants or microbes, is of great interest not only for researchers interested in human lung pathologies but also for environmental researchers using gills for toxicology studies and aquaculturists for whom preserving gill health is of paramount importance to ensure the stability of their stocks.

**Figure 1.**
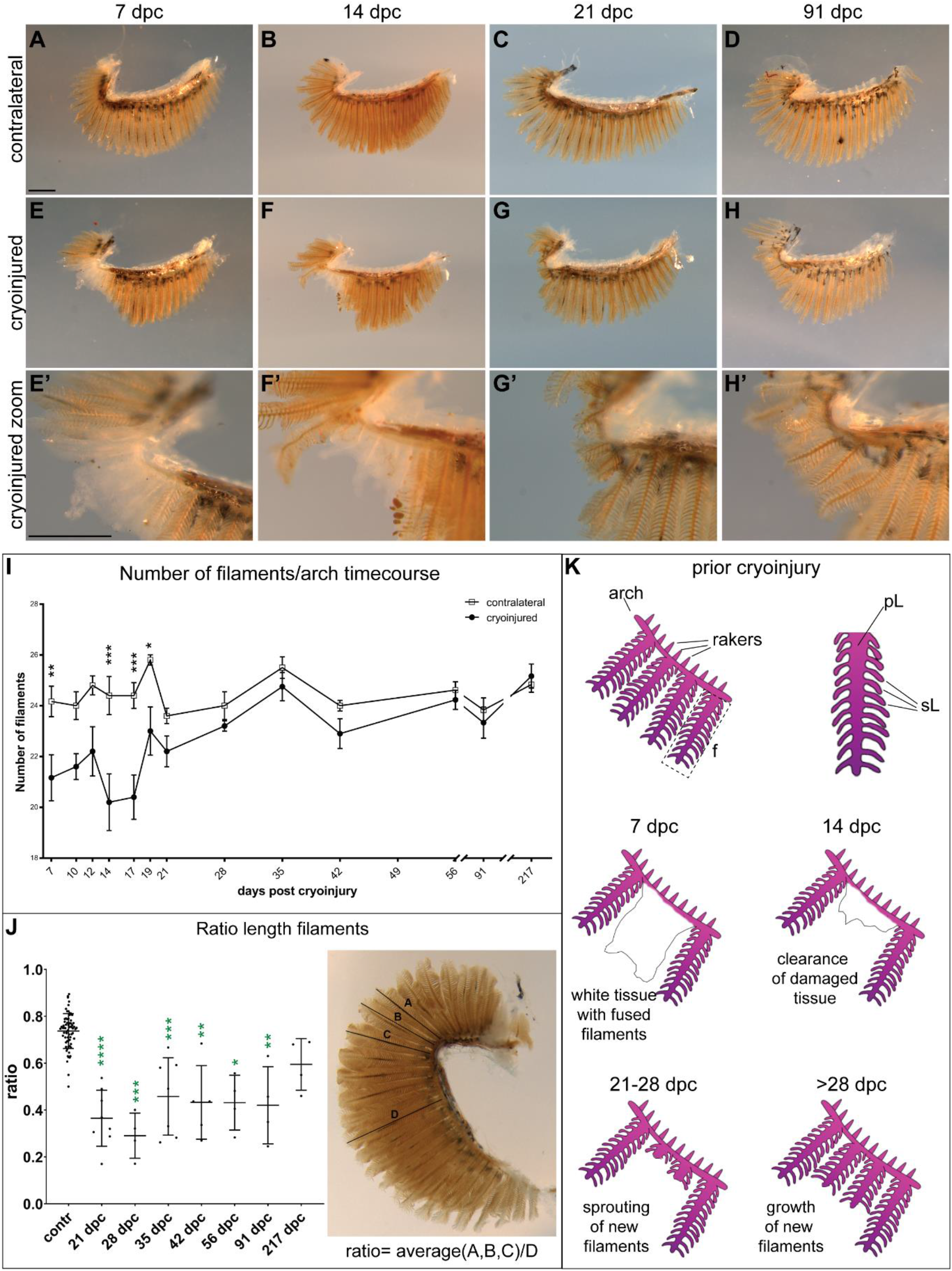
Gill filaments regenerate after cryoinjury. (A-H’) Brightfield images of dissected gill arches after cryoinjury. (E’-H’) Higher magnification images of the cryoinjured area in (E-H). In the examples shown, the contralateral and cryoinjured gills at each timepoint are from the same individual fish. Scale bar=0.5 mm. (I) Time course showing the number of filaments in dissected gill arches (contralateral and cryoinjured). Two-way ANOVA with Sidak’s multiple comparison test analysis was performed. Number of fish processed: n=6 (7, 91, and 217 dpc), n=5 (10, 12, 14, 17, 19 and 28 dpc), n=10 (21 and 42 dpc), n=8 (35 dpc), n=13 (56 dpc). (J) The growth of the regenerated filaments was evaluated using the ratio calculation as shown. One-way ANOVA test was used to compare individual cryoinjury timepoints with contralateral. n=69 for contralateral, n=8 (21 dpc), n=4 (28, 56, 91 and 217 dpc), n=7 (35 dpc), n=5 (42 dpc). Note that we could only measure the length of new filaments when these were straight and not overlapping each other. (I,J) *p<0.05, **p<0.01, **p<0.001; ****p<0.0001. (K) Graphical summary of the cryoinjury model. f=filaments, pL=primary lamella, sL=secondary lamella.

The zebrafish has an extraordinary ability to heal, repair, and regenerate damaged tissue not only in embryonic and larval stages but also at the adult stage (recently reviewed in [18]. To cause physical injury to zebrafish tissues, researchers have employed a variety of techniques such as resection and thermal wounding. Examples of tissue regeneration in zebrafish include the larval spinal cord and fin fold, the adult skin, heart, retina and caudal fin. In a recent study, Mierzwa et al (2020) showed that zebrafish gill filaments regenerate after resection. They found that resected filaments are 50% complete after ∼ 40 days and 85% complete after ∼160 days. The mechanism of gill filament regeneration after resection is believed to involve the formation of a blastema [12]. While very informative, this injury model could be further enhanced by studies that recapitulate the extensive tissue damage seen in chronic lung diseases. Cryoinjury of the zebrafish heart is suggested to be more representative of the cardiac tissue death observed in humans after ischaemia and myocardial infarction in comparison to simple resection models [19, 20]. This suggests that cryoinjury of zebrafish gills may similarly recapitulate at least some of the extensive and complex respiratory tissue damage observed in human lung tissue.

In this study, we developed a novel model of respiratory tissue damage involving cryoinjury of the adult zebrafish gills. We found that damaged gill tissue is able to repair and regenerate. Characterisation of the dynamics of repair and growth showed that immediately following injury, there is a period of tissue clearance with new filaments starting to grow 3 weeks after cryoinjury. An increase of both cytokine and chemokine transcripts with increased numbers of innate immune cells at the injury site was also observed. Analysis of various markers for apoptosis, proliferation, blastema formation and filament development suggest that there is a concurrent clearance of damaged tissue and tissue growth in cryoinjured gill tissue. RNA seq was used to investigate genome-wide transcript changes in the gills at different timepoints after cryoinjury allowing us to compare the transcriptional changes in our novel gill cryoinjury model with those of other zebrafish regeneration models and with those of mammalian respiratory tissue damage. Based on the transcriptional profiles, our findings suggest that there are mechanistic similarities but also differences in the way that various tissues repair damage and regenerate as well as similarities and differences when comparisons are made across species.

## Materials and Methods

### Zebrafish husbandry and fish lines

Zebrafish were maintained according to standard practice and all procedures conformed to UK Home Office requirements as per the Animals (Scientific Procedures) Act 1986. We used AB wild-type (WT) and the following transgenic lines: Tg(*lyz*:DsRed2) [21] and Tg(*KDR*:mCherry) [22]. The Tg(*mpeg1*:Caspase1 biosensor) transgenic line was generated in our laboratory using the published *mpeg1*.*1* promoter [23] and details of this line will be published elsewhere. For the purpose of this study, it is referred to as Tg(*mpeg1*:YFP). The transparent line used in this study was TraNac (*tra*^*b6*^ */tra*^*b6*^ ; *nac*^*w2*^ */nac*^*w2*^) [24], with some of the transgenic lines used either on a WT or TraNac background.

### Cryoinjury

A graphical schematic of the procedure is shown in Fig. S1. Adult fish were first anaesthetised in system water supplemented with MS-222 (168 mgs/mL; Sigma E10505). They were then placed in the lid of a petri dish with their right side facing up. Under a dissecting microscope, the operculum covering the right gill was removed using dissection scissors. Subsequently, the body of the fish and exposed gill arches were promptly dried with tissue paper, using the paper to absorb the moisture by capillarity. A steel probe (∼1 mm^2^ at point of contact) that was pre-cooled in liquid nitrogen was then applied to the exposed gill for 10 seconds before the fish was placed back in system water to recover. The left/contralateral gill arches were used as controls in our experiments. Fish were returned to the aquarium after the procedure. Analgesic measures were used during this procedure by dissolving Lidocaine Hydrochloride Monohydrate (Sigma L56647) in system water in the holding tank before the procedure (2.12 mg/L) and in the recovery tank after the procedure (2.65 mg/L).

### In situ hybridisation, staining, and histology

To generate *in situ* probes, partial or full cDNA fragments of zebrafish *sall4, cyclinb1*, and *fibina* were amplified from WT embryonic cDNA and cloned in pGEM®-T Easy (Promega). Primers used for each amplification are listed in Supplementary Methods (Table S1). For the *msx1b* probe, we used pCS2+-*msx1b* (gift from Bruce Riley). For the *mmp9* probe, we used pCRII-*mmp9* (gift from Anna Huttenlocher). All externally sourced and cloned plasmids were sequence verified. *in situ* hybridisation (ISH) was performed as described previously (Ramel and Hill, 2013). Synthesised digoxigenin-labelled probes were first tested on WT or TraNac embryos. Dissected gill tissue was fixed in 4% paraformaldehyde (PFA) overnight at 4°C, transferred to 1X PBS then processed for ISH, which included a 20 minutes proteinase K digestion (10 µg/mL) step at 37°C. Identical ISH results were observed using gill tissue from either WT or TraNac. Of note, ISH stained gills at times showed non-specific staining in the arch part of the gills and associated connective tissue. The Alcian Blue (cartilage) and Alizarin Red (bone) staining was adapted from previously published protocols [25, 26]. Immunostaining was essentially performed as described [9]. The following antibodies were used: Rabbit anti-mRFP antibody (1:1,000; MBL PM005), Rabbit anti-activated Caspase3 (1:500; BD Pharmingen 559565), Goat anti-Rabbit Alexa555 (1:300, ThermoFisher A32732) and Donkey anti-Rabbit Alexa555 antibody (1:500; abcam ab150074). Hoechst 33342 (1:1,000; Thermo Scientific™ 62249) or DRAQ5 (1:1,000; Thermo Scientific™ 62251) were used for nuclear staining. Histological sections of gills, Haematoxylin and Eosin (H&E) and Picrosirius Red (PSR) staining were processed as described [9].

### RNA seq and data analysis

Library preparation for sequencing was performed using 200 ngs of total RNA input with the TrueSeq RNA Sample Prep Kit v2-Set B (RS-122-2002, *Illumina Inc, San Diego, CA*) producing a 275bp Fragment including Adapters in average size. In the final step before sequencing, 8 individual libraries were normalised and pooled together using the adapter indices supplied by the manufacturer. Pooled libraries were clustered on the cBot Instrument (*Illumina Inc*) using the TruSeq SR Cluster Kit v3 - cBot – HS (GD-401-3001, *Illumina Inc*, San Diego, CA). Sequencing was performed as 50 bp, single reads and 7 bases index read on an Illumina HiSeq2000 instrument using the TruSeq SBS Kit HS-v3 (50-cycle, FC-401-3002, *Illumina Inc*). RNA seq data in FASTQ form from each sample were first quality assessed using fastQC. Reads were mapped using Bowtie [27] to the *Danio rerio* genome sequence v9 release 73 retrieved from Ensembl. Read counts for each exon and gene were generated in R using the GenomicRanges and Samtool Bioconductor packages against the *Danio rerio* v9 release 73 gtf file. Read counts were normalised to generate pseudo counts, followed by differential expression analysis using the edgeR [28] Bioconductor package which utilises empirical Bayes estimation and exact tests based on the negative binomial distribution. In addition, the Bioconductor package mmseq was used to obtain differentially expressed transcripts. Genes were deemed statistically differentially expressed using 0.01 as cutoff for the false discovery rate (fdr; Table S2).

### Comparative RNA seq analysis

Publicly available RNA seq datasets of adult zebrafish heart cryoinjury (GSE94617; [29], 3 dpc) and of adult caudal fin amputation (GSE76564, [30], 4 dpi) were chosen for comparative RNA seq of zebrafish regeneration models. Since these two datasets were generated using a pool of samples (one pool of 4 hearts and 2 pools of 10 caudal fins respectively) rather than individual samples, we selected the 1,000 genes with highest transcript levels in each model and dataset to allow a fair comparison (Table S3). This meant a log2(FC) >1.87 for the pools of heart and >1.85 for caudal fin. GO analysis was carried out using geneontology.org [31, 32] and Venn analysis was carried out using biovenn.nl [33]. For inter species comparison of respiratory tissue damage, the following RNA seq datasets were chosen: adult zebrafish cryoinjury gill (this study; 7 dpc), mouse lung bleomycin induced pulmonary fibrosis (PF) (Strobel et al., in preparation) (day 14), and mouse lung TGFβ1 induced PF [34] (day 14 and 21). Only genes with human orthologues were analysed. Note that we also verified some of the target genes identified in other published datasets (see Fig. 9).

### qPCR

Extraction of RNA and synthesis of cDNA for qPCR were essentially as described [9]. The Taqman gene expression assays used in this study are listed in Table S4. SYBR primers are listed in Table S5.

### Imaging

For imaging dissected gill arches, the heads of adult zebrafish were fixed in 4% PFA at 4°C for a maximum of 4 days. Gills were less fragile and easier to dissect using this procedure. After dissection, gill arches were placed in a drop of 1X PBS on a Petri dish containing 5% solidified agarose and imaged with reflected light using a Leica M205FCA stereomicroscope coupled with a Leica DFC7000 T camera. Images were acquired using the Leica LAS X imaging software. The Leica M205FCA microscope was also used to image Alizarin Red/Alcian Blue stained gill arches. For the mRFP immunostaining, images were acquired with an inverted Leica SP5 confocal microscope and the LAS AF software. For the activated Caspase 3 immunostaining, a widefield microscope was used (Zeiss Axio Observer 7) and images were processed using the Huygens deconvolution software. Images were further processed with FIJI (ImageJ), Adobe Photoshop and Adobe Illustrator to generate figures. N numbers in figure legends correspond to the number of gill arches imaged with the most representative images shown in the figure.

### Flow cytometry

Single cell suspensions of gill tissue from Tg(*lyz*:DsRed2) and Tg(*mpeg1*:YFP) adult zebrafish were obtained and processed for flow cytometry as previously described [9].

### Statistical analysis

Statistical analysis was performed using the Prism software (GraphPad Software, San Diego, CA). Unpaired two-tailed t-tests were used when comparing two groups. One-way or two-way ANOVA statistical tests were used for the analysis of time-course data (see Figure legends). p-values lower than 0.05 were considered to be statistically significant. *p<0.05, **p<0.01, ***p<0.001, ****p<0.0001.

## Results

### Zebrafish cryoinjured gill tissue is able to repair and regenerate within weeks

To cause physical damage to the adult zebrafish respiratory tissue, cryoinjury was carried out on one side of the adult gill by applying a pre-cooled metal probe (Fig. S1). To identify the different stages of gill tissue repair, we dissected and subjected to imaging the most external gill arches from both the cryoinjured and contralateral (uninjured control) sides at various times after cryoinjury (days post cryoinjury, dpc, Fig. 1A-H’). As early as 1 hour after cryoinjury, the affected filaments showed signs of damage with loss of their organised structure and detectable damage of primary and secondary lamellae. This was followed by fusion of the remaining gill filaments within the cryoinjured area into a tissue mass without any apparent organisation. The fused tissue mass persisted for at least 2 weeks (Fig. 1E-F). Interestingly, by 14 dpc the most distal part of this damaged tissue had been lost (Fig. 1F’). From about 3 weeks post-cryoinjury (21 dpc; Fig. 1G,G’), we observed that the damaged tissue was replaced by new budding gill filaments which continued to grow over time.

To quantify the changes observed, we counted the number of filaments (full length and partial length) of the most external arch on the cryoinjured side and the equivalent arch on the contralateral side of the same fish (Fig. S1; Fig. 1I). The fish used were 6-12 months old, and the number of filaments in contralateral arches ranged between 23 and 27. In the cryoinjured arches, due to the fusion of filaments after cryoinjury, there was initially a significantly lower number, between 17 and 25, of distinct filaments present. However, from 21 dpc we found that there was no significant difference in the number of filaments between contralateral and cryoinjured arches. This is consistent with our observation that new filaments start budding from ∼21 dpc (Fig. 1G). Of note, we found that these new filaments grew in length over time but did not fully re-grow as compared to the neighbouring non-cryoinjured filaments, even after 7 months (217 dpc, ∼81% of full length, Fig. 1J). Importantly however, new gill filaments appeared structurally normal and contained cartilage (Fig. S2) and blood vessels (Fig. S3) in which blood flow was observed (not shown).

These data show that after cryoinjury, the damaged gill tissue is progressively replaced, within a matter of weeks, by seemingly functional gill filaments which regrow but not quite to full length (summarised in Fig. 1K).

### Inflammation in the cryoinjured tissue is transient with innate immune cells recruited to site of cryoinjury in a temporally-regulated manner

Inflammation is prerequisite to initiate and successfully resolve tissue repair following injury [35-37]. Neutrophils and macrophages are rapidly recruited to the wound to allow clearance of damaged tissue, prevention of pathogen invasion and to provide signals (e.g. growth factors, metabolites) to other cell types such as fibroblasts. To characterise the inflammatory response following gill tissue damage caused by cryoinjury, we documented the immediate changes in the transcripts for pro-inflammatory factors as well as the accumulation of innate immune cells at the site of injury over several days.

We first used qRT-PCR to quantify the change in transcript levels of inflammatory cytokines and chemokines in gill tissue in the first 24 hours after cryoinjury (Fig. 2A-D). IL-1β is a cytokine produced by immune cells whose expression is increased after injury in zebrafish larvae [38]. *Il1β* transcript levels are also increased in the zebrafish gills after exposure to cigarette smoke or the viral adjuvant Resiquimod (R848) [9, 10]. After cryoinjury, *il1β* transcript levels rapidly increased with high levels at 9 hours post cryoinjury (hpc) and declining by 15 hpc (Fig. 2A). IL-6 is a cytokine that is produced by a variety of cells and its expression in zebrafish adults is increased after exposure to pro-inflammatory chemicals such as LPS and poly I:C [39]. *Il6* transcript levels are also increased in zebrafish gill tissue upon treatment with R848 [10]. As with *il1β*, we detected elevated levels of *il6* transcripts with high expression occurring at 15 hpc (Fig. 2B). Cxcl18b is a chemokine that attracts neutrophils upon injection of recombinant protein in the hindbrain and in a *Mycobaterium marinum* infection model in zebrafish larvae [40]. In the gills, *cxcl18b* transcripts increased rapidly at the site of cryoinjury (3 hpc) and remained at an elevated level over the duration of the time course (Fig. 2C). *Tnfα* transcript levels were also assessed but no consistent changes or differences were observed between cryoinjured and contralateral gill samples (Fig. 2D).

**Figure 2.**
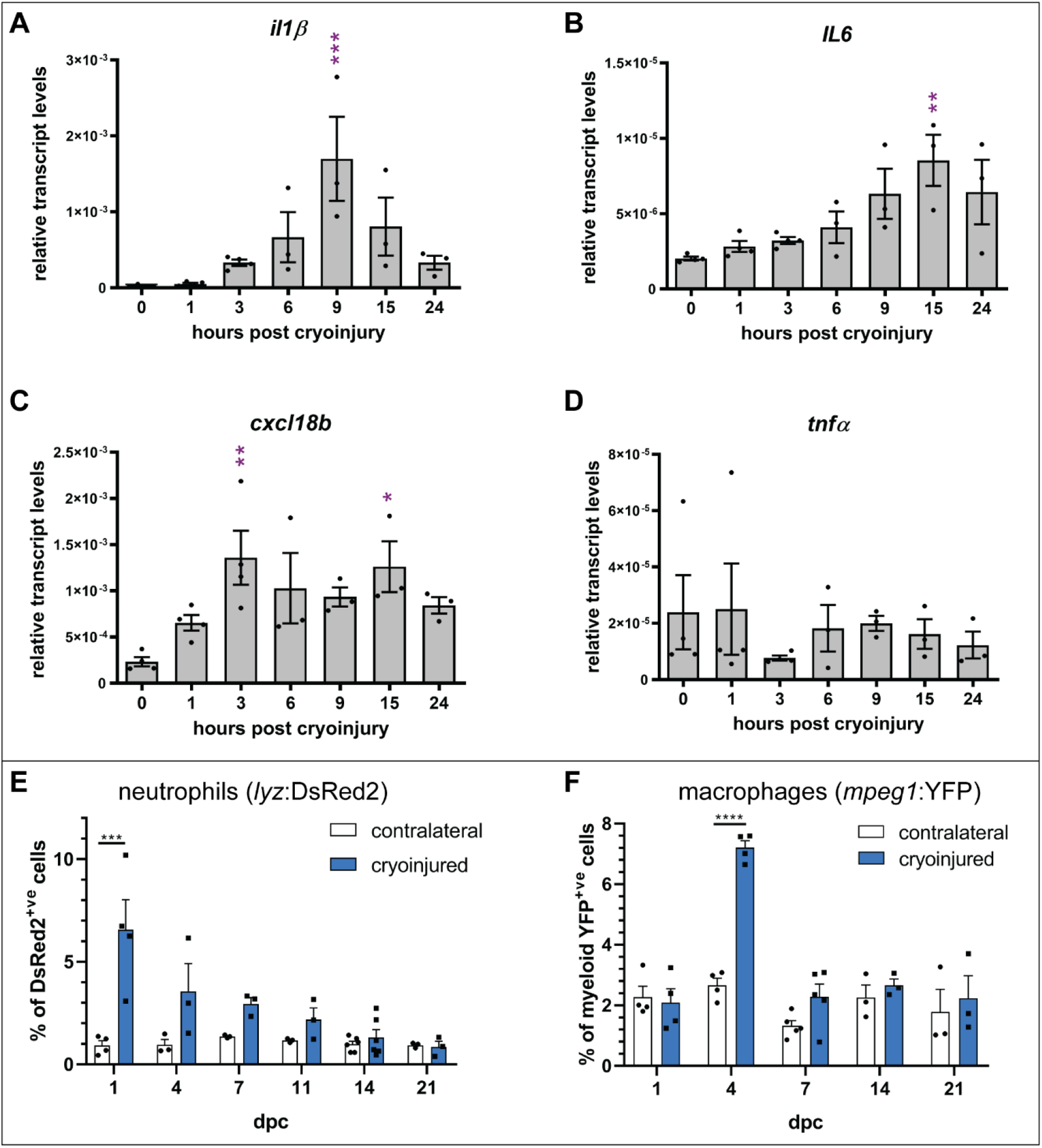
Transient inflammation occurs after gill cryoinjury. (A-D) qRT-PCR time course for *il1β, il6, cxcl18b, and tnfα* in the first 24 hours after cryoinjury. Transcripts were measured by qRT-PCR and are shown relative to 18S transcript levels. One-way ANOVA statistical test; significance is shown relative to 0 hours post cryoinjury. n=4 (0,1, and 3 hours post cryoinjury), n=3 (6, 9, 15, and 24 hours post cryoinjury). (E) Percentage of DsRed2^+ve^ cells (neutrophils) in cryoinjured vs contralateral gill tissue after flow cytometry analysis. n=4 (1 dpc), n=3 (4, 7, 11, and 21 dpc), n=6 (14 dpc). (F) As in E with percentage of YFP^+ve^ myeloid cells (macrophages). n=4 (1 and 4 dpc), n=5 (7 dpc), n=3 (14 and 21 dpc). (E-F) Two-way ANOVA statistical test. *p<0.05, **p<0.01, ***p<0.001.

After establishing an initial increase in cytokine and chemokine transcript levels, we investigated whether a cellular immune response could also be observed over the 3 weeks following cryoinjury. First, we made use of the zebrafish transgenic line *Tg*(*lyz*:DsRED2) [21] in which neutrophils are DsRed2 positive (DsRed2^+ve^) [41, 42]. Using flow cytometry, we observed an accumulation of DsRed2^+ve^ cells in cryoinjured tissue compared to the contralateral control side (Fig. 2E). The greatest accumulation of DsRed2^+ve^ cells was detected at 1 dpc and the number of cells decreased thereafter until 14 dpc when the percentages of DsRed2^+ve^ cells in cryoinjured and contralateral samples were similar. This result is consistent with the early induction of the neutrophil chemo-attractant Cxcl18b as detected by transcript levels (Fig. 2C). We then measured the dynamics of macrophage recruitment in cryoinjured tissue using our own *Tg*(*mpeg1*:YFP) transgenic line (see Materials and Methods). In other transgenic lines in which the same *mpeg1*.*1* promoter was used [23], labelled cells in adults belong to both the lymphoid and myeloid lineages, with the majority of myeloid cells being macrophages based on light scattering properties, single cell RNA-seq and cell morphology [43, 44]. We quantified the accumulation of YFP^+ve^ macrophages using flow cytometric analysis (Fig. 2F). We found that YFP^+ve^ macrophages are significantly more numerous at the cryoinjury site at 4 dpc compared to the contralateral side. At all other timepoints examined, levels of YFP^+ve^ cells are similar in contralateral and cryoinjured tissues.

To summarise, there is an inflammatory response in the zebrafish gill tissue after cryoinjury that is characterised by a rapid increase in pro-inflammatory transcripts. Following an early increase in neutrophils, macrophages accumulate at the site of damage. Importantly, the inflammatory response appears to be resolved by 2 weeks post injury.

### Collagen is deposited transiently at the cryoinjury site

While collagen deposition is necessary for tissue repair, prolonged deposition results in scarring and fibrotic changes. Following heart cryoinjury in the zebrafish, collagen is only transiently deposited [45-47]. To assess the level and duration of collagen deposition in the cryoinjured gills during the inflammatory stage (0-14 dpc) as well as during the growth phase, when filaments start growing back from 21 dpc, we performed histological analysis and qRT-PCR.

In gill tissue coronal sections stained with H&E (Fig. 3), we observed a complete fusion of filaments at 4, 7, and 14 dpc (Fig. 3B-D) with a noticeable increase in the amount of mucus cells at 4 dpc (arrows, Fig. 3B). At 21 dpc, consistent with the observations on dissected gills (Fig. 1), distinct filaments began to be visible (Fig. 3E) and by 42 dpc, the gross gill architecture was restored (Fig. 3F). To visualise the dynamics of collagen deposition, we also stained the sections with Picrosirius Red (PSR) and used polarised light to identify the collagen-rich structures. In normal, contralateral gills, collagen is detected in the filaments (arrows, Fig. 3A’). Collagen deposition is altered at 4 and 7 dpc (Fig. 3B’-C’) with a visible increase in PSR staining detected at 7 dpc (Fig. 3C’). From 14 dpc onwards (Fig. 3D’-F’), the normal pattern of collagen appears to be restored. We performed qRT-PCR for *col1a1a* which encodes a structural protein that is a constituent of collagen type I. Consistent with the PSR staining data, there was a significant increase in *col1a1a* transcript levels at 7 dpc in cryoinjured gill tissue when compared to the contralateral gill (Fig. 3G).

**Figure 3.**
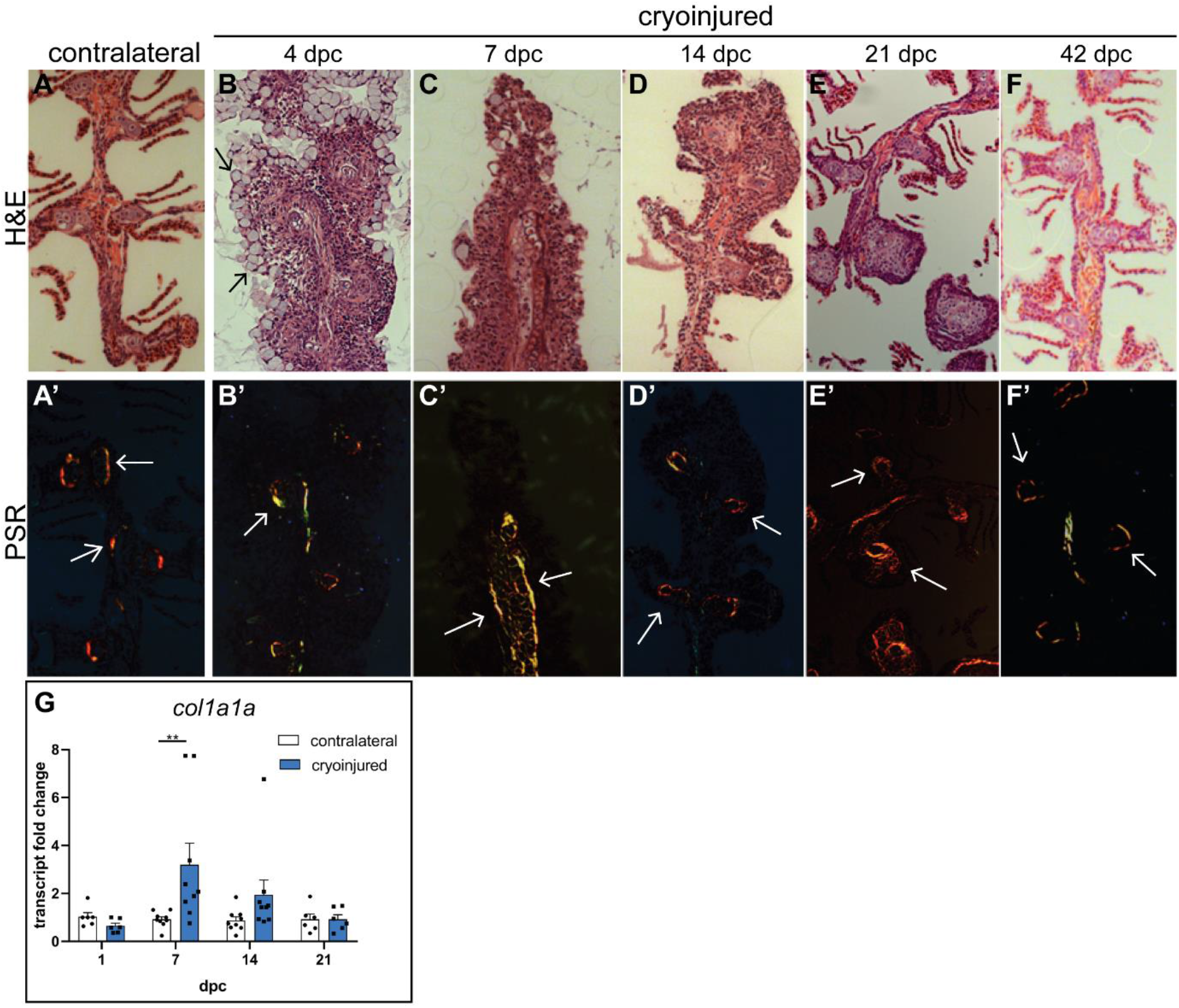
Filament fusion and transient collagen deposition are detected in cryoinjured gills. (A-F) H&E staining of gill coronal sections. Arrows in (B) indicate mucus cells. (B-D) No distinct filaments are seen. (A’-F’) PSR staining of gill coronal sections. Collagen-rich structure are indicated by white arrows. (G) qRT-PCR time course of *col1a1a* expression in contralateral and cryoinjured gill tissue (normalised to 18S, results are expressed as fold change). Two-way ANOVA test: **p<0.01. n=6 (1 and 21 dpc), n=9 (7 dpc and 14 dpc).

In summary, increased collagen deposition occurs transiently following cryoinjury with the maximum observed being at 7 dpc. No persistent abnormal collagen deposition was detected in the damaged tissue, highlighting the ability of the zebrafish to repair gross physical wounding to a respiratory tissue without scarring.

### Cellular apoptosis and proliferation patterns indicate an active process of regrowth within cryoinjured gill tissue

Our results indicate that for the first 2 weeks the cryoinjured tissue is a hub for inflammation and collagen deposition (Figs 2,3). This is followed by a period of tissue clearance and new tissue growth (Fig. 1). Indeed, after the resolution of inflammation (from 2 weeks post injury), the damaged tissue is replaced by new filaments that extend distally over time (Fig. 1). To further understand the cellular events during this complex tissue repair process, we characterised the patterns of cellular apoptosis and proliferation in the cryoinjured tissue.

We used activated Caspase 3 immunofluorescence [48] to identify apoptotic cells in zebrafish gills at 3 timepoints, namely 6 hpc, 3 dpc, and 7 dpc. In contralateral gills, there appeared to be a small number of apoptotic cells located throughout the gill arches, mostly along the filaments (Fig. 4 A-C). At all timepoints analysed we observed an increase in the number of apoptotic cells in the cryoinjured area compared to both surrounding non-cryoinjured gill tissue and contralateral tissue (Fig. 4D-F’). Interestingly, we also observed increased apoptosis in the filaments just adjacent to the cryoinjured area (Fig. 4D’-F’); this could be because these filaments were partially damaged by the cryoinjury process. Overall, the activated Caspase3 immunostaining results are suggestive of an increase in cellular death in the first week after cryoinjury which can explain the loss of damaged tissue observed after 7 dpc (Fig. 1).

**Figure 4.**
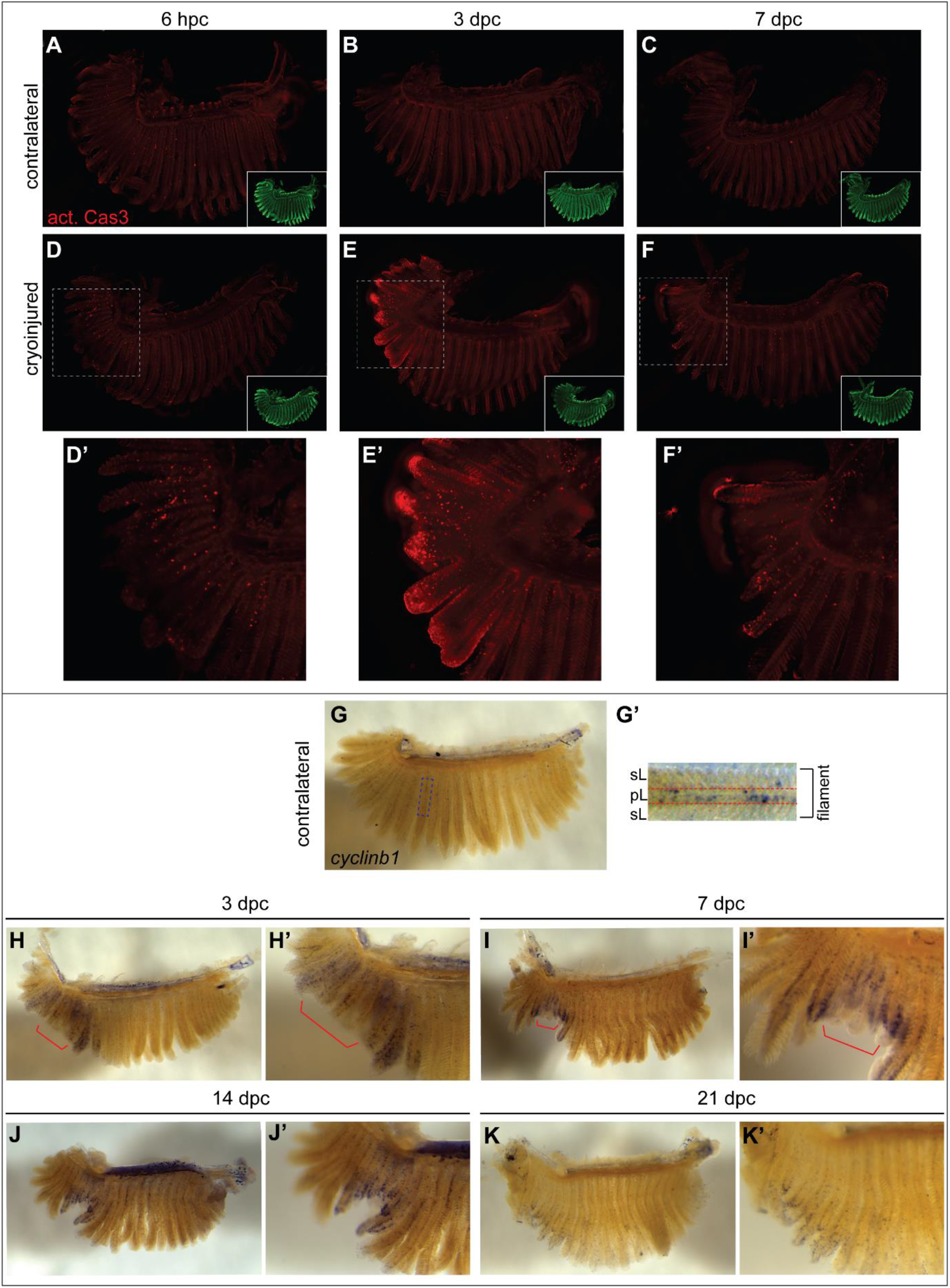
Apoptosis and proliferation in cryoinjured gill tissue. (A-F) Activated Caspase 3 immunofluorescence in contralateral (A-C) or cryoinjured gill arches (D-F) at 6 hpc (n=4 contralateral, n=6 cryoinjured), 3 dpc (n=4 contralateral, n=6 cryoinjured), and 7 dpc (n=3 contralateral, n=3 cryoinjured). Dashed squares (D-F) indicate the cryoinjured area. (D’-F’) Magnified views of the cryoinjured area outlined in (D-F). Images in the bottom right corner of (A-F) are the corresponding DRAQ5 nuclear staining. *Disclosure: gill images were acquired in a different orientation (∼90*^*o*^ *C angle) so the corners of each image were peudocoloured black to generate panels A-F*. (G-K’) ISH for *cyclinb1* in contralateral and cryoinjured gill arches. G’ is a magnified image of the dashed rectangle in G with pL=primary lamella and sL=secondary lamellae. H’, I’, J’, and K’ are zoomed in versions of H, I, J, and K respectively. Red brackets (H-I’) highlight the central area of the cryoinjured tissue. *cyclinb1* ISH: 3 dpc (n=5 contralateral, n=7 cryoinjured), 7 dpc (n=4 contralateral, n=4 cryoinjured), 14 dpc (n=4 contralateral, n=6 cryoinjured), 21 dpc (n=7 contralateral, n=5 cryoinjured). TraNac fish were used in these experiments.

To document the proliferation patterns in cryoinjured gill tissue, we carried out *in situ* hybridisation (ISH) for *cyclinb1*. Cyclinb1 is a cyclin that regulates the G2-M transition during the cell cycle and the *cyclinb1* mRNA expression pattern is an indicator of cellular proliferation in the adult zebrafish retina [49]. In contralateral gills, low levels of *cyclinb1* were detected along the primary lamellae, at the base of secondary lamellae (Fig 4G-G’). This pattern is consistent with the expression of the proliferation markers PCNA and BrdU/idU in the zebrafish and Medaka gills [11, 50, 51]. At 3 and 7 dpc, many proliferating cells were detected in the cryoinjured area (Fig. 4H-I’). However, the most central part of the cryoinjured area (red brackets, Fig. 4H-I’) appeared to contain many fewer proliferating cells compared to the filaments immediately adjacent to that central area. This suggests that the most affected cells in the centre of the cryoinjured tissue, where significant apoptosis was detected (Fig. 4A-F’), are dying and not dividing at 3 and 7 dpc. The *cyclinb1* expression pattern was different at 14 and 21 dpc with a large number of proliferating cells present in the distal half of the cryoinjured tissue and affected filaments (Fig. 4J-K’); this expression pattern is restricted to the tips/apical domains of the growing filaments at 21 dpc (Fig. 4K-K’). In teleost gills, cells in the apical domain of filaments proliferate to allow growth and addition of new secondary lamellae [50, 51], making the location of *cyclinb1* expressing cells at 21 dpc consistent with the growth of newly regenerated filaments observed at that stage (Fig. 1).

To summarise, analysis of apoptosis in the gill cryoinjured tissue showed that a large number of cells undergo apoptosis in the first week after cryoinjury. This occurs synchronously with an increased proliferation in the filaments directly adjacent to the most badly damaged central area of the cryoinjured tissue. Subsequently, the proliferation pattern indicates that cells in the most distal parts of the filaments are dividing to drive apical growth.

### Regeneration and growth of zebrafish gill filaments occurs via activation of developmental programmes

To identify the regeneration mechanisms that drive growth of new filaments in cryoinjured gills the expression of markers associated with blastema formation and gill filament development was investigated.

Msx1b (muscle segment homeobox 1b) is a homeodomain transcription factor whose expression is associated with blastema formation after resection or cryoinjury of the zebrafish caudal fin [52-54]. ISH for *msx1b* in zebrafish gills indicated that it is not expressed in normal, contralateral gills (Fig. 5A). However, *msx1b* transcripts were readily detectable in cryoinjured gill tissue at 7 and 14 dpc (Fig. 5B,C). At these timepoints, expression of *msx1b* was observed primarily in the middle of the damaged tissue (arrows, Fig. 5B’,C’) which corresponds to the mass of fused filaments that eventually is shed (Fig. 1). *msx1b* expression at 21 dpc was much weaker with faint expression detected at the apical end of growing filaments (arrow, Fig. 5D’). The *msx1b* expression pattern suggests that there is a period after cryoinjury (7-14 dpc) where blastema formation is concurrent with inflammation, collagen deposition, apoptosis and proliferation.

**Figure 5.**
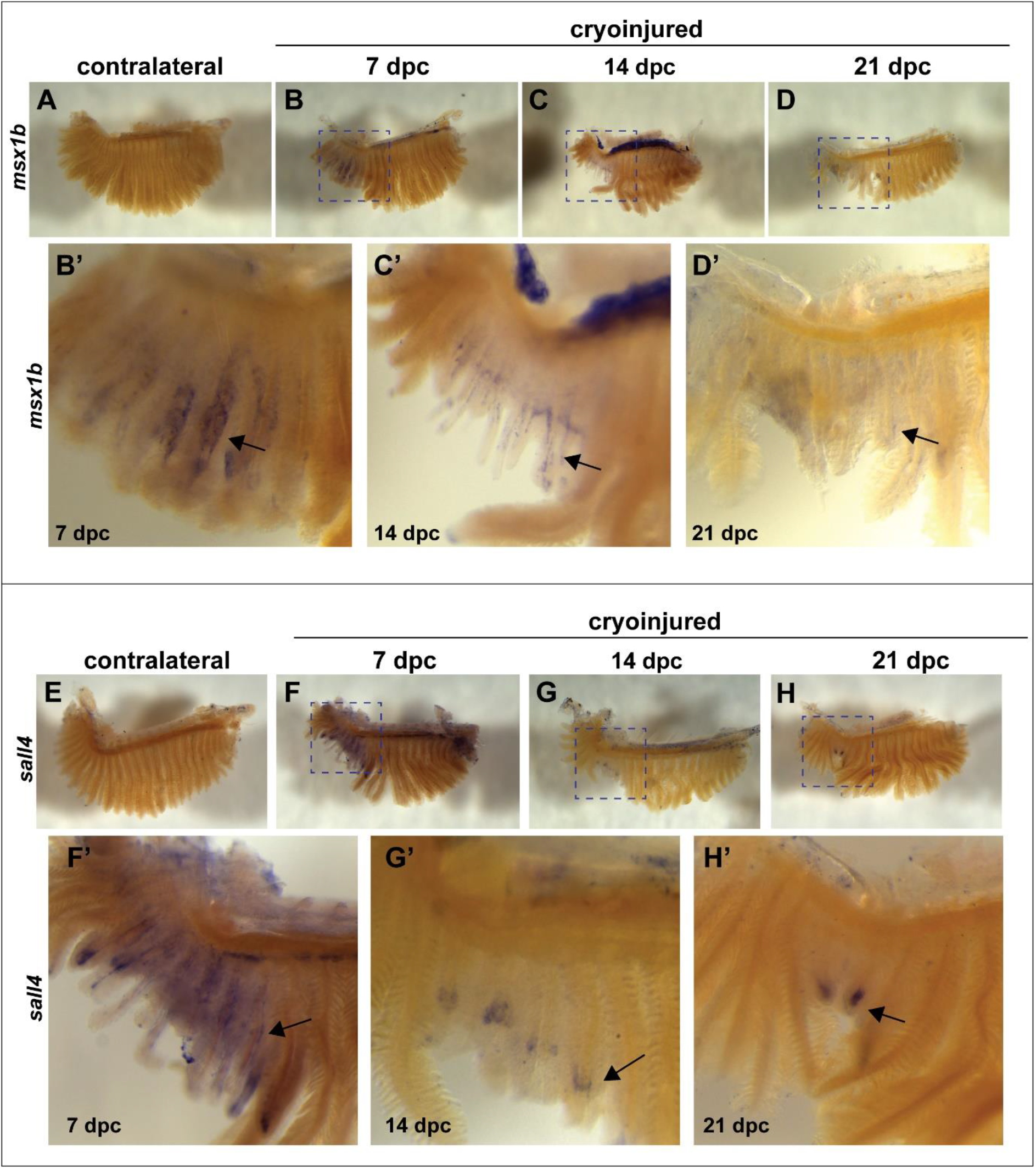
*msx1b* and *sall4* are expressed dynamically in cryoinjured gill tissue. (A-D’) ISH for *msx1b* in contralateral (A) and cryoinjured gills (B-D). (B’-D’) Magnified views of cryoinjured areas (corresponding to dashed squares in B-D). Arrows (B’-D’) point to *msx1b* mRNA deposition. Only faint expression is seen at 21 dpc (D’). Contralateral (n=26), cryoinjured 7 dpc (n=9), 14 dpc (n=7), 21 dpc (n=12). (E-H’) ISH for *sall4* in contralateral (E) and cryoinjured gills (F-H). (F’-H’) Magnified views of cryoinjured areas (corresponding to dashed squares in F-H). Arrows (F’-H’) point to *sall4* mRNA deposition. *sall4* is detected at most distal tips of growing filaments at 21 dpc (H’). Contralateral (n=28), cryoinjured 7 dpc (n=14), 14 dpc (n=14), 21 dpc (n=6). TraNac fish were used in these experiments.

Sall4 (spalt-like transcription factor 4) is a transcription factor known to regulate embryonic stem cell pluripotency and self-renewal [55]. In zebrafish, *sall4* is a pluripotency marker that is enriched in blastomeres [56] and in 10 days old zebrafish, *sall4* mRNA is also specifically enriched at the tip of the gill filament primordia, suggesting that it could be important for their development [57]. The expression of *sall4* in cryoinjured gills was investigated using ISH and transcripts were not detected in normal contralateral gills (Fig. 5E). It was however expressed in the cryoinjured gill area at all timepoints examined (Fig. 5F-H). Interestingly, its expression also became restricted to the tip of regenerated filaments at 21 dpc (arrow, Fig. 5H’) which suggests that Sall4 may contribute to the growth of filaments observed after 21 dpc.

*msx1b* and *sall4* expression in cryoinjured gills suggests that a blastema-like tissue is formed and may contribute to the initiation of a developmental program for filament growth and differentiation.

### Transcriptomic analysis of gill cryoinjury

To investigate the underlying mechanisms that drive wound healing and repair we performed transcriptomic analysis using bulk RNA seq of gill tissues at 3, 7, 14 and 28 dpc (Fig. 6A).

**Figure 6.**
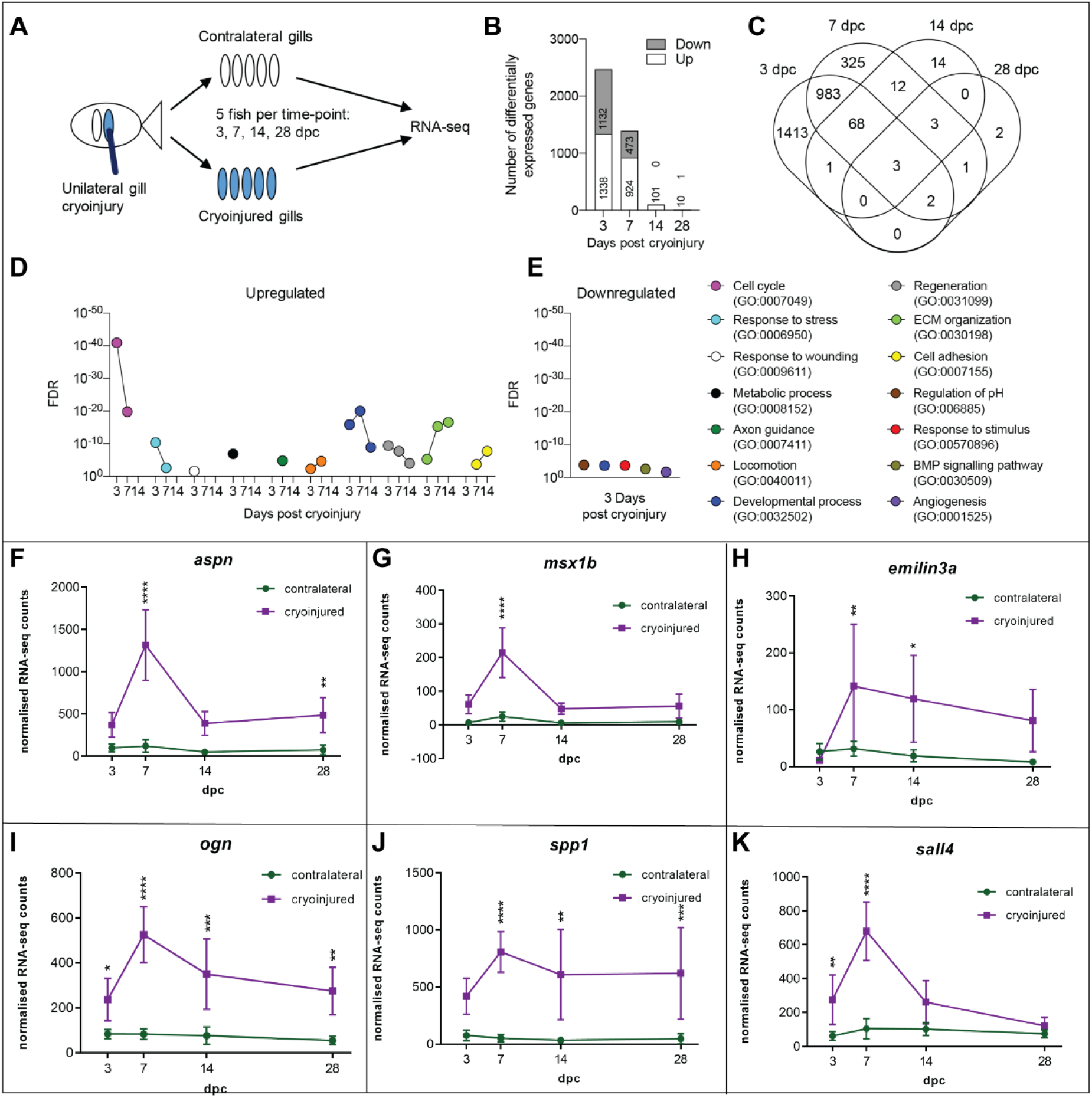
RNA seq time course analysis of cryoinjured gill tissue. (A) Schematic illustrating the samples submitted for RNA seq (see also Fig. S1). (B) Number of differentially expressed genes across time. (C) Venn diagram showing the number of differentially expressed genes that are shared between time-points. (D-E) Selected GO terms for increased and decreased transcript levels at indicated time-points. (F-K) Normalised RNA seq counts in contralateral and cryoinjured samples for *aspn* (F), *msx1b* (G), *emilin3a* (H), *ogn* (I), *spp1* (J), and *sall4* (K) at 3, 7, 14, and 28 dpc. Two-way ANOVA statistical test: *p<0.05, **p<0.01, ***p<0.001, ****p<0.0001.

Comparative analysis of the cryoinjured area and the contralateral side revealed that the number of differentially expressed genes was highest at 3 dpc and gradually decreased thereafter (Fig. 6B; Table S2). Overlap analysis showed that some genes were differentially expressed at specific time points only, while others appeared across a longer timeframe suggesting an activation of stage-specific transcriptional programmes (Fig. 6C). To interrogate the biological processes taking place at the various time-points we performed gene ontology (GO) analysis of the differentially increased and decreased gene transcripts (Table S2). In agreement with the data shown in Fig. 4, 3 dpc was dominated by GO terms related to cell cycle (Fig. 6D). In addition, injury-related GO terms such as response to stress and response to wounding were present at 3 dpc. GO terms associated with immune responses were absent from 3 dpc and all other time-points investigated. Indeed, in contrast with qRT-PCR results from the first 24 hours after cryoinjury where changes in inflammatory markers were detected (Fig. 2), analysis of inflammatory cell marker genes (*mpx, mpeg1, lyz, lcp1, cd4* and *lck*; Fig. S4A-F) and pro-inflammatory cytokines such as *tnfβ, tnfα*, and *il1β* (Fig. S4G-I) showed that these were indeed not altered upon cryoinjury during the analysed timeframe from 3 dpc onwards. Consistent with results shown in Fig. 2, *csf1ra* (a macrophage marker gene) showed a slight increase increase at 3 dpc although not significant (Fig. S4J). Taken together, these results indicate that the acute pro-inflammatory phase following cryoinjury is transient and has mostly resolved by 3 dpc. Metabolic process-related GO terms were abundant amongst genes whose transcripts were increased at 3 dpc, while genes whose transcripts were decreased displayed GO terms for regulation of pH, developmental process, response to stimulus, BMP signalling pathway and angiogenesis (Fig. 6E) reflecting the complex responses following cryoinjury-induced tissue damage of the lamellar respiratory capillary system. 7 dpc was dominated by cell cycle-related GO terms, albeit less significantly compared to day 3, as well as terms related to developmental process, extracellular matrix organisation and regeneration. Consistent with ISH results (Fig. 5), the most significant increase in *msx1b* and *sall4* transcripts occurred at 7 dpc (Fig. 6G,K) Interestingly, we found an abundance of nervous system-related GO terms such as axon guidance at 7 dpc suggesting that gill innervations regenerate during that time-point. At 14 dpc, GO terms such as extracellular matrix (ECM), cell adhesion and locomotion were abundant highlighting processes such as new tissue formation and tissue remodelling. We found no significant GO terms associated with the 28 dpc time-point, however transcripts of genes associated with extracellular matrix and tissue formation such as *aspn, emilin3a* and *ogn* were sustained (Fig. 6F,H-I). The most increased gene at 28 dpc was *spp1*/*osteopontin* (Fig. 6J), which has previously been associated with zebrafish heart tissue repair and collagen type I deposition [58].

### Comparative transcriptomic analysis between zebrafish gills, heart and caudal fin following injury

In order to compare the gill regeneration program with other well characterised zebrafish injury models, we performed comparative transcriptomic analysis of our data set at 3 dpc with RNA seq datasets of the heart at 3 dpc (GSE94617, [29]) and the caudal fin at 4 days after transection (GSE76564, [30]). Overlap analysis of the top 1,000 genes whose transcripts were increased (Table S3) showed that there was only about 15% of genes shared between all three injury models, while there were more genes expressed in both cryoinjured gills and caudal fin cut (332) when compared to cryoinjured gills and heart (236; Fig. 7A). Interestingly however, overlap analysis of GO terms highlighted that biological processes were mostly shared between the three injury settings, again with a greater overlap between cryoinjured gills and caudal fin cut when compared to gills and heart (Fig. 7B). Common processes shared by all models with a similar significance were terms related to cell division (Fig. 7C). The most striking difference between the three models was the continued presence of immune response-related GO terms in the heart at 3 dpc, while these were completely absent from both the gill and the caudal fin. GO terms related to tissue remodelling such as developmental processes, ECM organisation, regeneration and cell adhesion, were shared between all three models however their expression indicated again a greater similarity in the processes involved in gill and caudal fin injury repair than in the heart and either gill or caudal fin.

**Figure 7.**
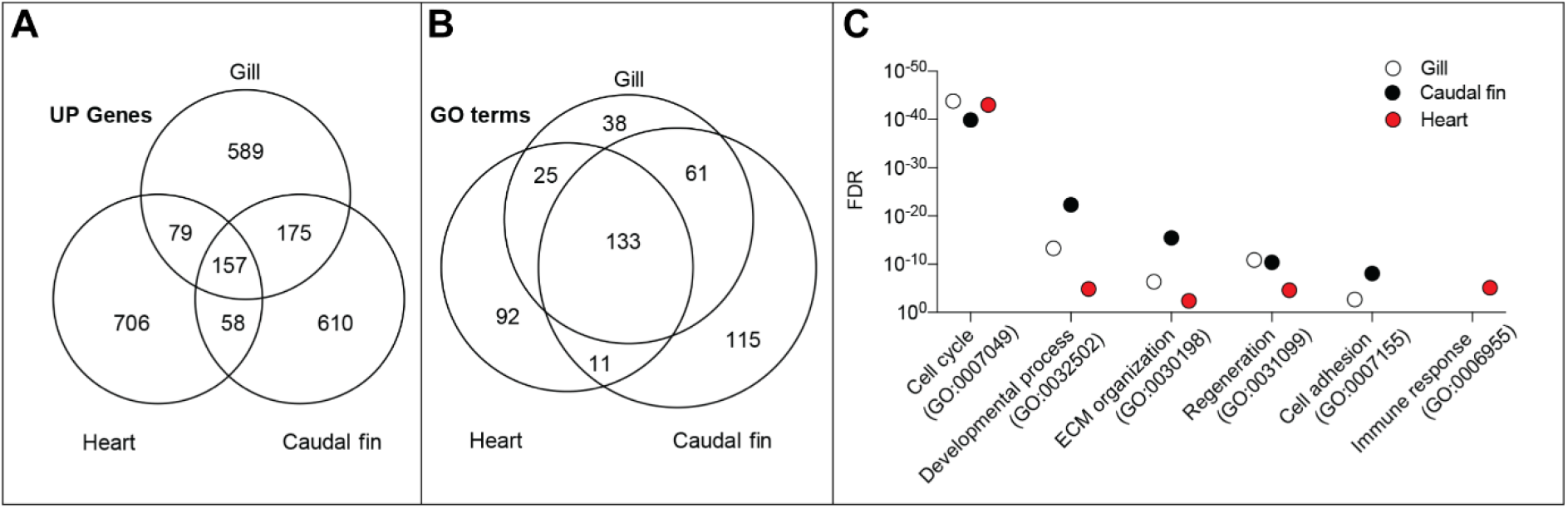
Comparative gene ontology analysis for gill cryoinjury, heart cryoinjury and caudal fin amputation. (A-C) Venn diagrams showing (A) the top 1,000 upregulated genes and (B) the gene ontology terms between gill cryoinjury, heart cryoinjury, and caudal fin amputation. (C) Selected gene ontology terms and their associated FDR for each injury condition.

Our comparative transcriptomic analysis of 3 different adult tissue injury models in zebrafish at a similar timepoint suggests that the initiation of repair programs in the gill and the caudal fin are more similar than in the heart after cryoinjury. Indeed, we have shown that the regeneration of the gill filaments appears to be epimorphic as observed in caudal fins (see Fig. 5; [59]).

### Comparative transcriptomic analysis of gill cryoinjured tissue and mouse models of pulmonary fibrosis

We have shown that the adult zebrafish respiratory tissue is able to repair and regenerate without scarring after substantial damage. In contrast, repair of the human distal lung is variable, with fibrous scar tissue progressively replacing healthy tissue after substantial/continuous tissue damage in conditions such as Idiopathic Pulmonary Fibrosis [60, 61]. To identify the genetic/transcriptomic basis for these differential repair abilities, we compared the transcriptome of cryoinjured zebrafish gills at 7 dpc (peak of expression of ECM remodelling genes and regeneration factors; see Fig. 6) and the transcriptome of mammalian models of lung fibrosis at peak fibrosis timepoints: mouse lung Bleomycin treatment model at day 14 (Strobel et al., in preparation) and mouse lung TGFβ1 overexpression model at day 14 and 21 [34]. Only differentially expressed genes in the respective datasets that have human orthologues were selected for this comparison. We focused on genes whose transcripts were inversely differentially expressed in zebrafish vs mouse to identify potential genes/pathways that account for the different tissue repair properties (Fig. 8A,C). Among the transcripts increased in mouse pulmonary fibrosis (PF) models and decreased in our zebrafish gill cryoinjury model (Fig. 8A) are the chemokine CXCL14, the chemokine receptor ACKR3, and the ubiquitin ligase RNF128. This is suggestive of some persistent inflammation in mouse PF models at the chosen timepoints while inflammation in cryoinjured zebrafish gills at 7 dpc is already past its peak (see Fig. 2). We confirmed via qRT-PCR that zebrafish *rnf128a* transcripts are decreased in cryoinjured gills when compared to contralateral gills, not only at 7 dpc, but also at 3 dpc (Fig. 8B). We also identified a number of genes whose transcripts were increased in zebrafish cryoinjured gills while decreased in mouse PF models (Fig. 8C). Interestingly, a number of genes that were identified are known to be involved in fibroblast biology and extracellular matrix remodelling: FGFBP1, ITGA8, MMP15, and MMP9. Notably, we identified FIBIN (Fin Bud Initiation Factor) as a gene whose transcripts are differentially expressed in our comparative analysis; with transcripts of both zebrafish paralogues (*fibina* and *fibinb*) increased in cryoinjured gills while Fibin transcripts are decreased in mouse PF models. We confirmed via qRT-PCR the increase in zebrafish *mmp9* (Fig. 8D), *fibina* (Fig. 8E), *fibinb* and *gelsolinb* (not shown) transcripts in cryoinjured gills. We also used ISH to visualise the increase of *mmp9* and *fibina* transcripts in the gill cryoinjured area at 7 dpc (Fig. 8F-K). ISH results clearly showed that *mmp9* and *fibina* are not normally detected in gill tissue (Fig. 8F,I) while they were both strongly expressed in the cryoinjured tissue (Fig. 8G-H,J-K). To sum up, our inter species comparative analysis not only highlighted some of the mechanistic differences between the two models (such as transient vs persistent inflammation and ECM remodelling) but also identified potential targets of interest.

**Figure 8.**
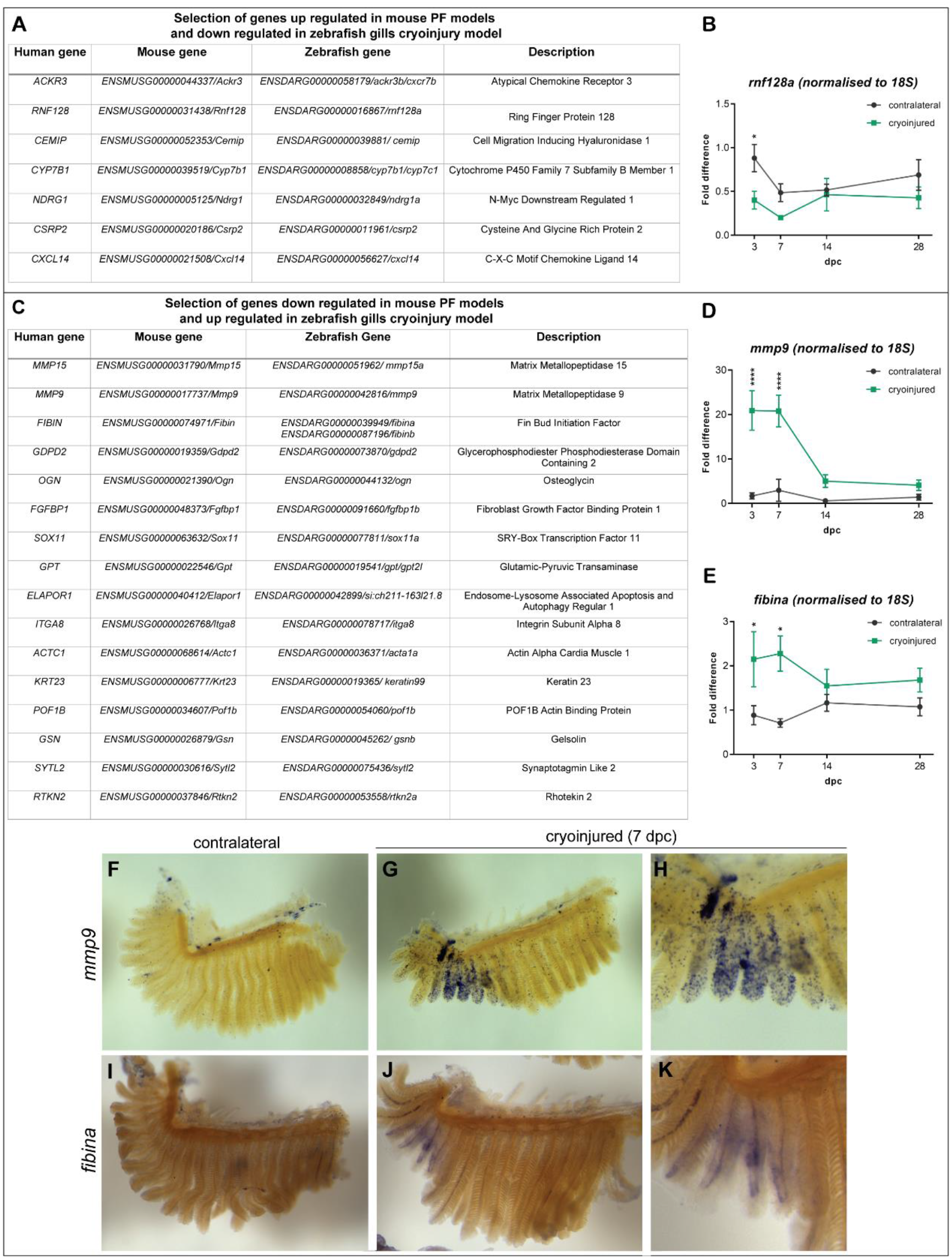
Comparative RNA seq analysis. (A) List of genes whose transcripts were increased in mouse PF models and decreased in zebrafish gill cryoinjury model. (B) qRT-PCR analysis of zebrafish *rnf128a* after gill cryoinjury. (C) List of genes whose transcripts were decreased in mouse PF models and increased in zebrafish gill cryoinjury model. (D-E) qRT-PCR of zebrafish *mmp9* (D) and *fibina* (E) after gill cryoinjury. (F-H) ISH for *mmp9* in contralateral (F) and cryoinjured 7 dpc gill arch (G, higher magnification in H). (I-K) ISH for *fibina* in contrateral (I) and cryoinjured 7 dpc gill arch (J, higher magnification in K). (B,D,E) Two-way ANOVA statistical test * p<0.05, **** p<0.0001.

**Figure 9.**
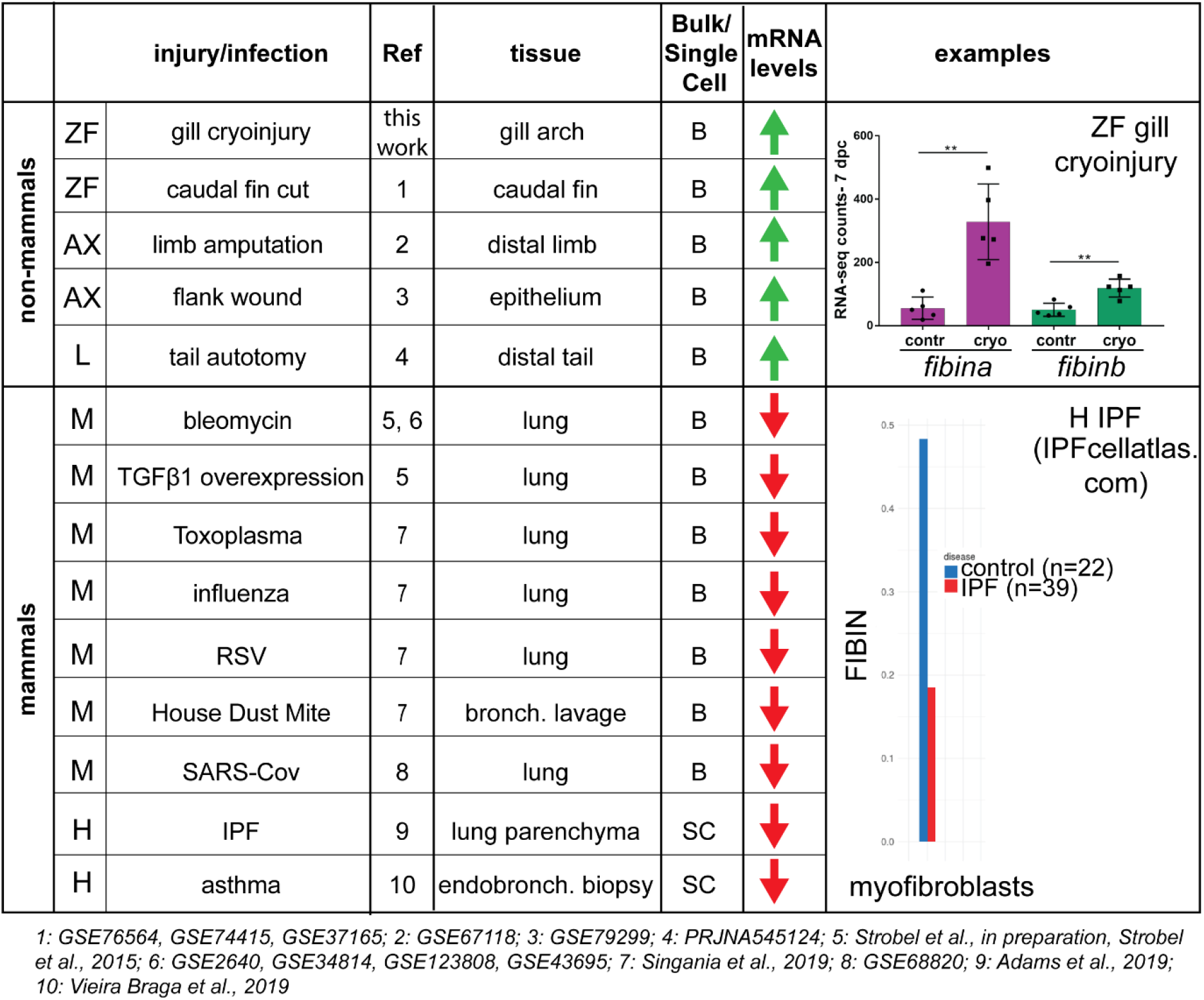
Data mining of FIBIN expression levels in non-mammalian models of tissue damage and mammalian models of respiratory tissue damage or infection. ZF=zebrafish; AX=axolotl; L=lizard; M=mouse; H=human. Material used for transcriptomic analysis: bulk tissue (B) and single cells (SC). The numbers of the datasets analysed are indicated as well as article references which are linked to specific online search databases [73-75] Examples of FIBIN mRNA levels in zebrafish gills (contralateral vs cryoinjured) and human lung myofibroblasts (control vs IPF) are shown in the last column.

### Datamining of Fibin expression in various models of injury and repair

Our RNA seq comparative analysis identified FIBIN as a gene whose transcripts are increased during zebrafish gill regeneration (*fibina* and *fibinb*) while mouse *Fibin* transcripts are decreased in models of pulmonary fibrosis. Very little is known about FIBIN function either in zebrafish or mouse. The only functional study to date showed that zebrafish Fibinb is necessary for pectoral fin bud development (morpholino study; [62]). Protein prediction analysis tools failed to predict Fibin function with confidence (not shown). There is also conflicting evidence that Fibin is a secreted protein [62, 63]. Using publicly available transcriptomics datasets, we investigated the expression of FIBIN in various models of respiratory tissue damage, as well as repair and regeneration models in different organisms (Fig. 9, including references and NCBI Geo datasets numbers). Strikingly, we found that *fibin* transcripts are increased in several non-mammalian models of tissue regeneration. Indeed, not only are zebrafish *fibina* and *fibinb* transcripts increased in our gill cryoinjury model, they are also increased after caudal fin amputation. *fibin* transcripts are also increased during repair of flank wound in axolotl and during tail regrowth in the lizard. In contrast, in mouse models of respiratory tissue damage such as Bleomycin treatment and TGFβ1 expression overexpression, *Fibin* transcripts are down regulated. *Fibin* transcripts are also decreased in lung samples of mice exposed to house dust mite, Toxoplasma, or *influenza*, RSV, and SARS-Cov viruses. Analysis of single cell RNA seq datasets in zebrafish (caudal fin; [64]) and the human lung (IPFcellatlas.com) also revealed that zebrafish *fibina, fibinb* and human FIBIN are expressed in the mesenchymal lineage, specifically in fibroblasts (not shown). Consistent with the expression of FIBIN in lung fibroblasts and with the decrease of *Fibin* transcripts in mouse models of lung damage, we found that mesenchymal FIBIN expression is decreased in lung samples from patients with chronic lung conditions such as IPF and asthma (Fig. 9). Our data mining investigations thus uncovered the dichotomy of FIBIN expression in organisms well known for their regenerative capacity in comparison to mammalian species with induced or chronic respiratory tissue damage. FIBIN may therefore have a role in respiratory tissue repair and the regulation of its expression may be key to the differential repair abilities of different organisms.

### Summary of results and discussion

We have developed and characterised a novel model of respiratory tissue injury in zebrafish. Cryoinjury of gill tissue resulted in a transient inflammatory response followed by concomitant collagen deposition, apoptosis, cellular proliferation and the initiation of developmental programmes to promote filament growth and tissue regeneration.

### Concurrent events in cryoinjured gills

Cryoinjury is a complex type of injury compared to sterile cut models where the different stages of immune response, healing, and regeneration have been well defined. This is because cryoinjury causes progressive loss of tissue and progressive growth of new tissue concurrently. Overlap of different phases of repair have also been reported in a zebrafish caudal fin cryoinjury model [52] with the exception that no collagen deposition was detected in this model. Concurrent tissue growth and transient fibrotic scarring have also been observed in the zebrafish heart after cryoinjury [46, 65].

### Inflammatory immune response after cryoinjury

We have shown that there is a rapid induction of cytokine/chemokine transcripts in cryoinjured gill tissue that is accompanied by an accumulation of neutrophils and then macrophages at the site of injury. This sequence of events is similar in other adult zebrafish regeneration models such as the heart [65, 66] and caudal fin [67]. Resolution of inflammation appeared to occur by 14 dpc based on the flow cytometric analysis of neutrophil and macrophage accumulation. Our RNA seq analysis of inflammatory markers also confirmed that there is no sustained chronic inflammation in the cryoinjured gill tissue. As inflammation resolution is a prerequisite for tissue repair [35], the dynamics of inflammation activation/de-activation in our model is favourable for tissue repair. The contribution of macrophages to gill tissue repair is of interest as macrophages can have a dual function (pro-inflammatory vs pro-regenerative). Indeed, in zebrafish, pro-regenerative *wt1b*^+ve^ macrophages have been identified that may be important for timely repair of heart and caudal fin tissue after injury [68]. In the human lung, airway macrophages are critical in maintaining tissue homeostasis and controlling inflammation to allow repair of damaged lung tissue (reviewed in [69]). In light of this, the investigation of macrophages in gill tissue repair in our novel injury model could inform the mechanisms at play in the human lung.

### Extent of regeneration of zebrafish gills after cryoinjury

After cryoinjury, we found that new filaments start growing from 3 weeks. However, even after 7 months, regenerated filaments, which to all intent and purposes appear morphologically normal, did not reach the length of neighbouring uninjured filaments or filaments from contralateral gills. This would imply that this zebrafish respiratory tissue only achieves partial regeneration, as opposed to the fin fold and caudal fin for instance. In their study, Mierzwa et al. (2020) also reported that regenerated gill filaments fail to reach full length after resection. The authors suggested that this is because gills continue growing during adulthood as the fish get bigger and longer. Indeed, gills must grow to guarantee oxygen supply as adult fish keep growing. A recent study in Medaka also highlighted the continuous growth of gills over adulthood and identified two adult stem cell pools in the gills: branchial arch stem cells that generate extra filaments on the lateral edges of the gill arches and filament stem cells that contribute to the apical growth of filaments [50]. These stem cells allow for continuous growth of the gills during adulthood. In our model, we suggest that newly regenerated filaments have to grow in a tissue that is growing itself, which means that they are not quite able to ‘catch up’ their length to that of neighbouring filaments.

### Regenerated filaments are morphologically normal

Despite their smaller length, it appeared that regenerated filaments were morphologically normal (gross organisation, cartilage, vascularisation, blood flow). Investigating the expression of various markers for all the gill cell types (e.g. neurones, pavement cells, goblet type cells) will be key to fully assert that regenerated filaments are completely normal at the cellular level. Regarding functionality, being able to accurately measure gas exchange in regenerated gill filaments represents a technical challenge. As regenerated filaments do not reach full length, this means that the number of secondary lamellae (respiratory surface) is lower in those filaments compared to uninjured ones. This would imply that gas exchange may be slightly different in contralateral compared to repaired cryoinjured gills. The zebrafish heart cryoinjury model was developed 10 years ago [45-47] and at the time all studies pointed to complete regeneration of the heart and myocardium after cryoinjury. However, recent studies using more advanced technologies showed that zebrafish hearts are not completely regenerated and functional after cryoinjury [19]. Future work in our gill cryoinjury model may be able to address the full functionality of regenerated gill tissue.

### Conservation of transcriptional programmes for repair and regeneration

Comparative RNA-seq analysis of our gill cryoinjury dataset with published heart cryoinjury and caudal fin amputation datasets suggests that there is closer similarity in the gene ontology class of genes whose transcripts were increased within damaged gills and caudal fin compared to within gills and heart. The GO terms in common between gills and caudal fins are for developmental processes, ECM organisation and regeneration. The regeneration of limbs in neonatal mice, axolotl and salamander and the regeneration of the zebrafish caudal fin are known to involve the formation of a blastema and the redeployment of embryonic developmental programs for regeneration [59, 70, 71]. The basic development and growth of gill filaments can be reminiscent of appendages such as fins and limbs: gill filaments are protrusions which use apical growth to extend distally [50]. The tip of a zebrafish gill filament has actually been likened to a plant meristem with apical cells driving outgrowth to generate new secondary lamellae [51]. The fact that a known blastema marker, *msx1b*, is expressed in cryoinjured tissue also supports the theory of conserved repair mechanisms between gill filaments and fins. It is therefore not surprising that some of the mechanisms of regeneration of gills and caudal fins would be similar. In fact, a recent study showed that gill arches and fins in the skate develop from a common pool of precursor cells (neural crest and mesoderm) [72], so these structures may have retained common developmental (and regeneration) programmes. In addition to the similarities and differences observed in terms of gene ontology, our work also highlighted that each tissue has its own distinct set of repair and regeneration transcriptional instructions. We believe that our novel model provides a great platform to research the molecular mechanisms of vertebrate respiratory tissue repair. Indeed, comparative transcriptome analysis between our zebrafish gill cryoinjury model versus mouse models of lung fibrosis has identified genes of interest, including FIBIN. Datamining of FIBIN expression in various injury and repair scenarios clearly showed the power of comparative RNA seq analysis to identify target genes worth of pursuing. We believe such an approach could lead the way to the identification of new respiratory tissue repair pathways and to the development of future therapies.

## Acknowledgements

We thank Boehringer Ingelheim for funding. We are grateful for the core facilities in the Imperial College Life Sciences department (Central Biological Services, SAFB Flow Cytometry Facility, Facility for Imaging by Light Microscopy) and for the help provided by Lorraine Lawrence for the histological work. We would like to thank Tobias Hildebrandt, Werner Rust, and Germán Lepar for performing RNA seq as well as Karsten Quast, E. Simon, Geraint Barton and the Imperial College Bioinformatics Data Science Group for help with the analysis. Thanks to the following researchers for providing lines and/or reagents: Paul Martin, Steve Wilson, Bruce Riley, Anna Huttenlocher, and Steve Renshaw. Finally, we would like to thank Terrence Cook, Dory Polos, David Miao, Evan Kont, and past and present members of the Dallman laboratory for useful discussion and support.

