## Supplemental data for "Dynamics of repair and regeneration of adult zebrafish respiratory gill tissue after cryoinjury"

**Supplementary Materials and Methods**

**Table S1**

List of primers used to clone *cyclinb1, sall4*, and *fibina.*

| Construct | Forward primer | Reverse primer |
|---|---|---|
| pGEM®-T Easy-*cyclinb1* | cgagtcacagcaataaaccacgag | ttcccagtaacttcctttcctgc |
| pGEM®-T Easy-*sall4* | acggtgcagagtcagatgaa | acccctggagattgtggaag |
| pGEM®-T Easy-*fibina* | atgtggacgtcgcctctgtc | ttaaacgttcagatagtcgt |

**Table S2**

Excel file. Normalised RNA-seq counts for differentially expressed genes at 3, 7, 14 and 28 dpc. See Materials and Methods for details.

**Table S3**

Excel file. Gene Ontology comparative analysis of the top 1,000 up regulated genes in cryoinjured gills (3 dpc), cryoinjured heart (3 dpc), and amputated caudal fin (4 dpi). See Materials and Methods for details.

**Table S4**

List of Taqman gene expression assays used in this study.

| Target gene | Applied Biosystems assay number/order number |
|---|---|
| *18S* | 4310893E |
| *il1β* | Dr03114368_m1 |
| *il6* | Custom ARGZGHF PN444114 |
| *tnfα* | Dr03126850_m1 |
| *cxcl18b* | Dr03436643_m1 |
| *col1a1a* | Dr03150834_m1 |
| *rnf128a* | Dr03146167_m1 |
| *mmp9* | Dr03139882_m1 |

**Table S5**

List of SYBR primers used in this study.

| Target gene | Forward primer | Reverse primer |
|---|---|---|
| *fibina* | acctttgctccccgaaatgt | tctgcggtcgatccatttga |

**Supplementary figures and legends.**

**Figure S1**

Graphical outline of the cryoinjury procedure.

The right gills of adult zebrafish are exposed and cryoinjured using a steel cryoprobe. The gills on the left side of the fish (contralateral) are untouched and used as control. At specific timepoints after cryoinjury, the gill arches are dissected. Full arches are used for imaging and for counting filaments. The cryoinjured areas of the gills arches are further dissected out and processed for transcriptomics or flow cytometric analysis. The equivalent areas on the contralateral side are also collected to serve as control.

Figure S1

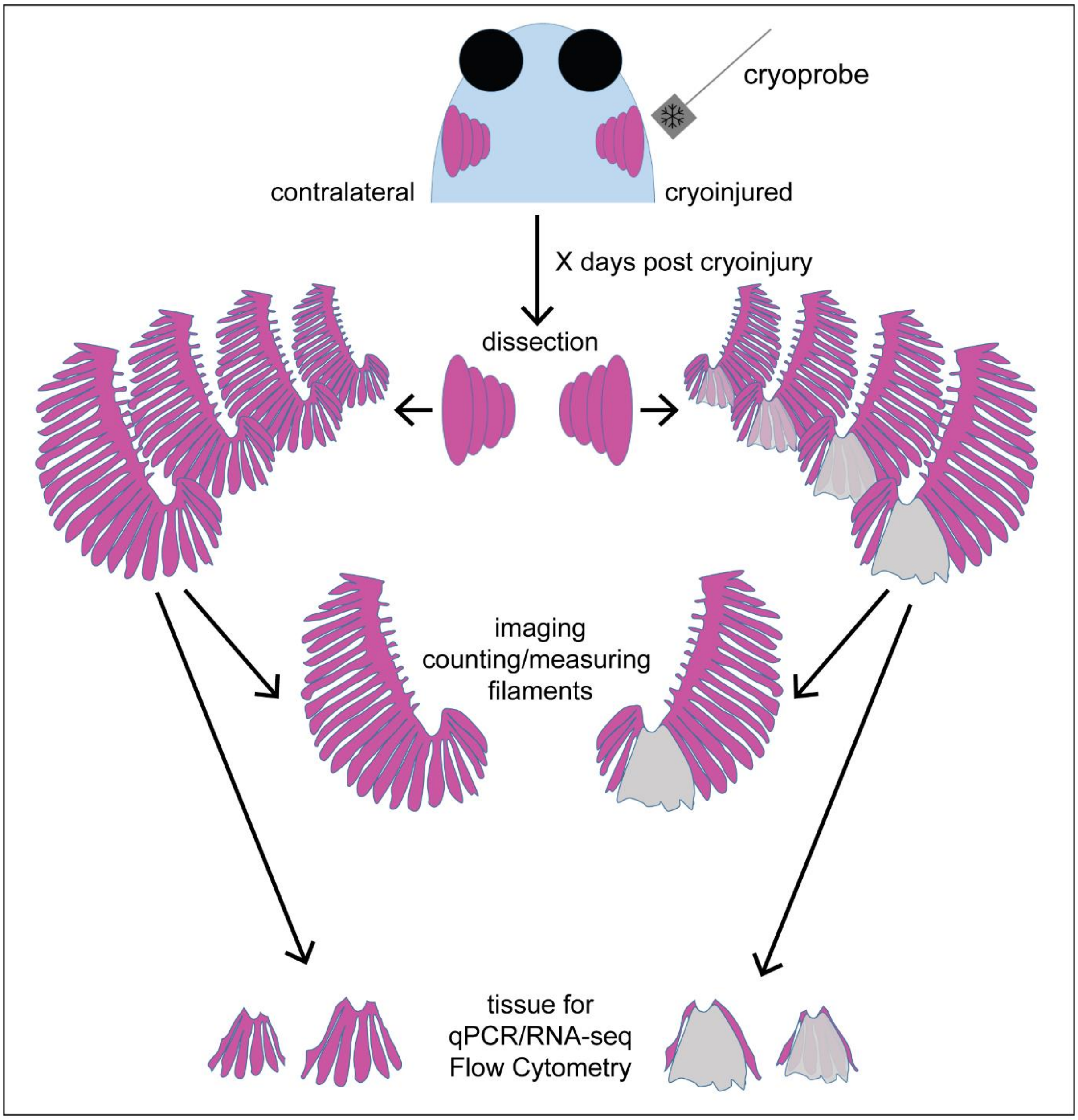
cryoprobe
contralateral
cryoinjured
X days post cryoinjury
dissection
imaging
counting/measuring
filaments
tissue for
qPCR/RNA-seq
Flow Cytometry

**Figure S2**

Newly regenerated gill filaments contain cartilage.

(A-B') Alizarin Red and Alcian blue staining of contralateral and cryoinjured gill arches. 28 dpc. (B') shows the area in the dashed rectangle in (B) with new filaments staining positive for Alcian blue. n=3 contralateral, n=5 cryoinjured. TraNac fish were used in this experiment.

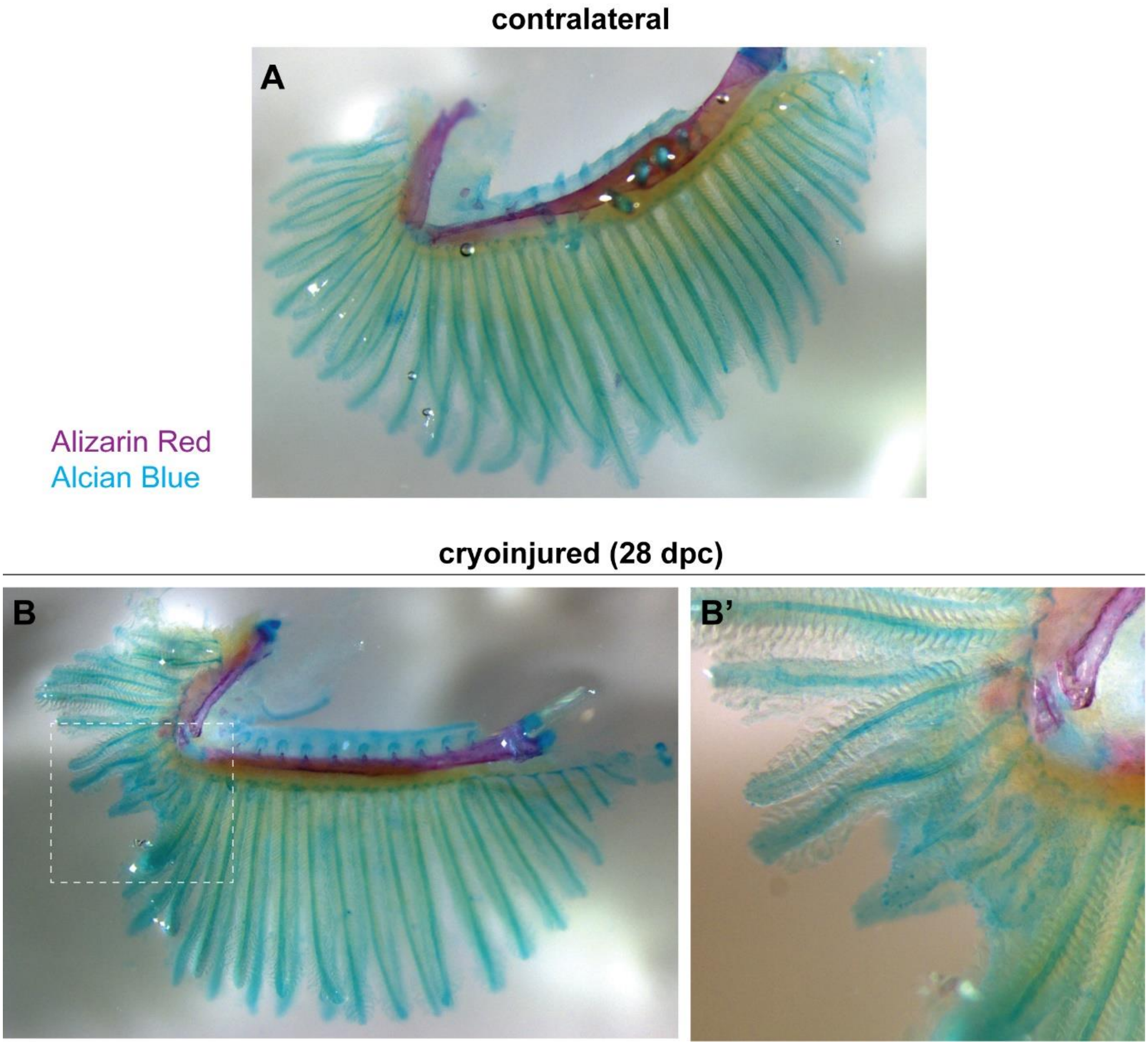

**Figure S3**

Newly regenerated gill filaments are vascularised.

Cryoinjured gill arch of TraNac Tg(*KDR*:mCherry) (42 dpc) immunostained with anti-mRFP antibody (B,B’; pseudocoloured magenta). Hoechst is used for nuclear staining (A, A’; pseudocoloured grey). The new filament highlighted in the dashed boxed in A is vascularised (B,B’). Scale bar=100 µm.

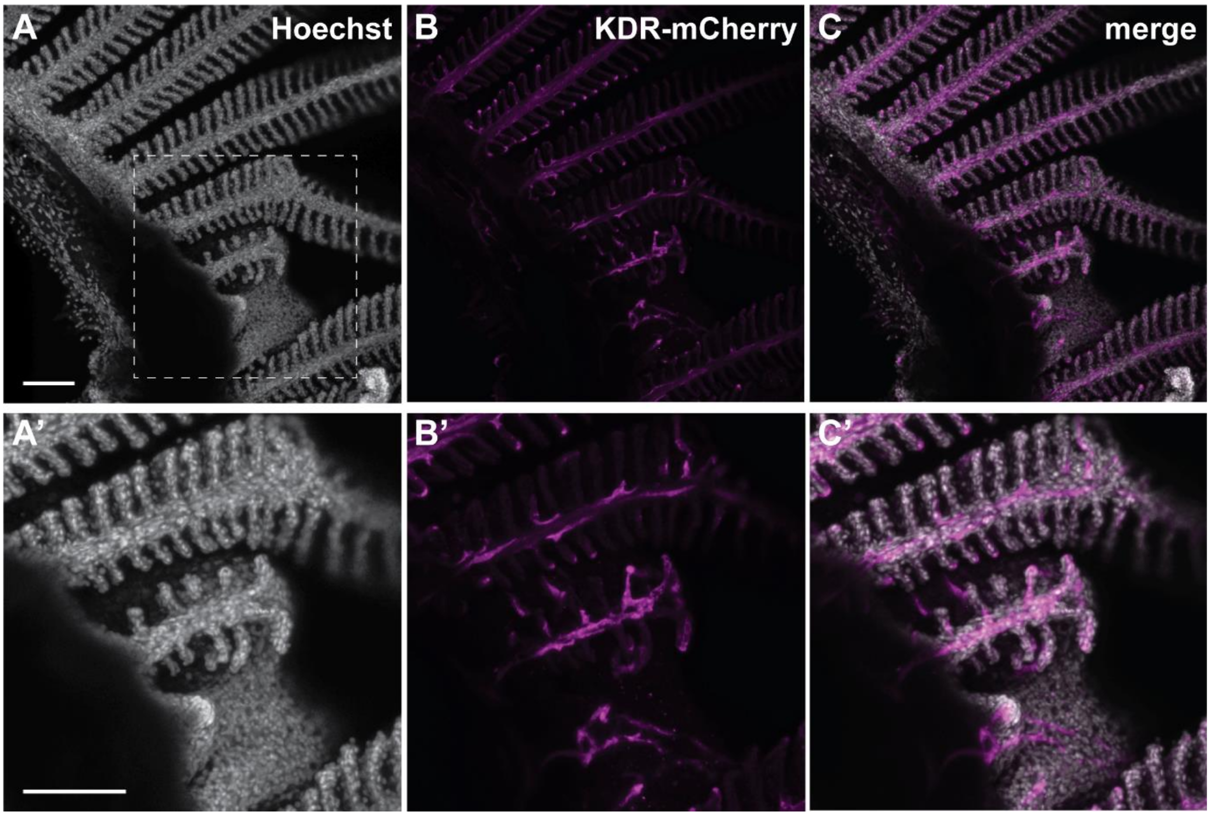

**Figure S4**

RNA seq analysis of inflammatory markers after gill cryoinjury.

Normalised RNA-seq counts for *mpx* (A), *mpeg1* (B), *lyz* (C), *lcp1* (D), *cd4* (E), *lck* (F), *tnfβ* (G), *tnfα* (H), *il1β* (I) and *csf1ra* (J) at 3, 7, 14, and 28 dpc. Error bars are not displayed when they are shorter than the height of the symbol. Two-way ANOVA statistical test: no significant differences.

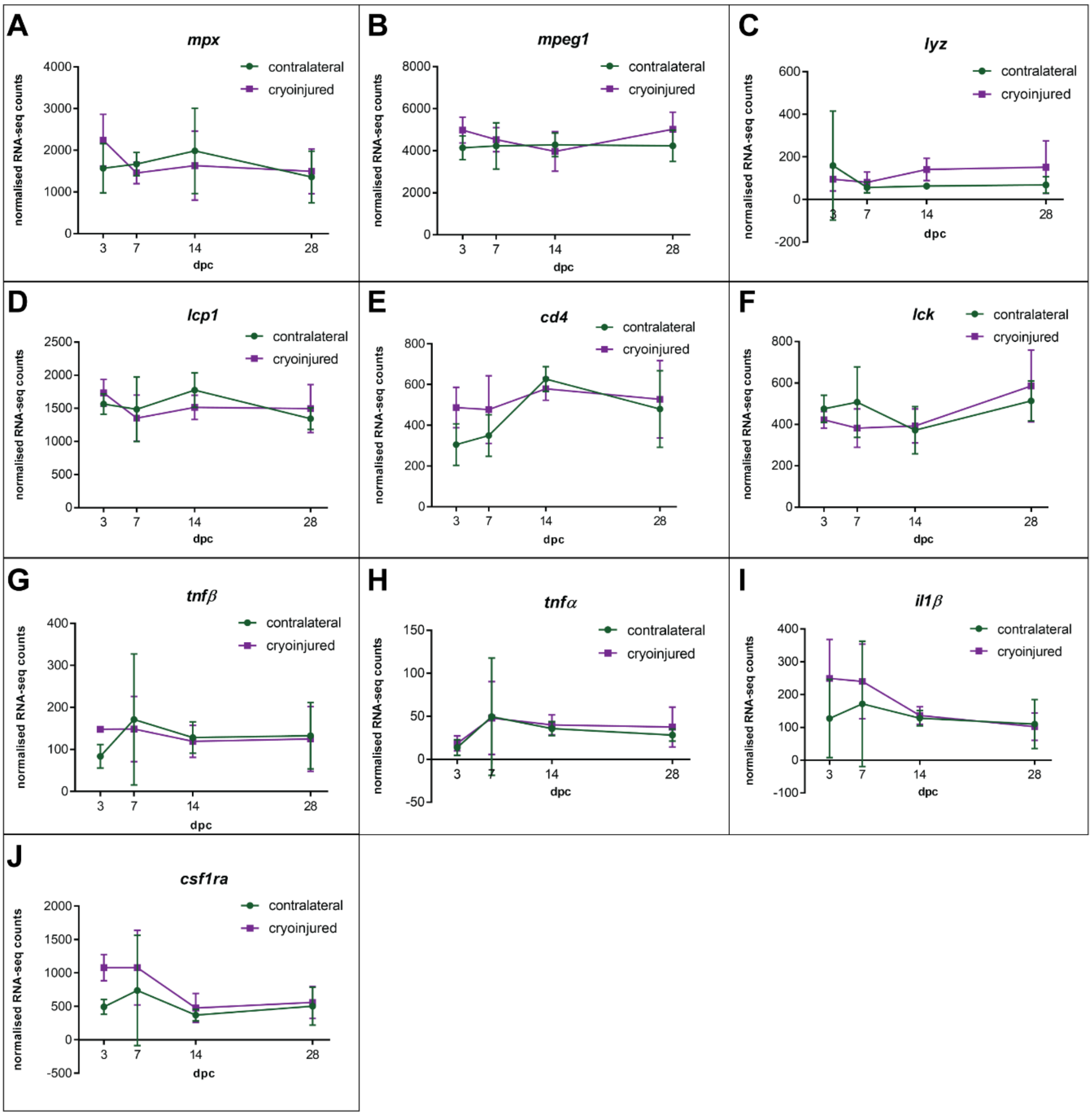
